# Cooperation with non-kin: Context-dependent acceptance of alien queens by polygynous ant workers

**DOI:** 10.1101/2020.11.27.401513

**Authors:** Ornela De Gasperin, Pierre Blacher, Michel Chapuisat

**Author notes:** Correspondence:Pierre Blacher, Department of Ecology and Evolution, University of Lausanne, 1015 Lausanne, Switzerland. These authors contributed equally to this work (Ornela De Gasperin, Pierre Blacher).

## Abstract

Relatedness underlies the evolution of reproductive altruism, yet eusocial insect colonies occasionally accept unrelated reproductive queens. To better understand this seemingly paradox, we investigated whether acceptance of unrelated queens by workers is an incidental phenomenon resulting from failure to recognize non-nestmate queens, or whether it is an adaptive behavior favored in specific contexts where cooperation is preferable to rejection. Our study system is the socially polymorphic Alpine silver ant, *Formica selysi*. Within populations some colonies have a single queen (monogynous), and others have multiple, sometimes unrelated, breeding queens (polygynous). Social organization is determined by a supergene with two haplotypes. In a first experiment we investigated whether workers from polygynous colonies were inherently more prone to accepting unrelated queens than workers from the alternate, monogynous social form. We found that workers rejected all alien queens, independently of their social origin and of the number of queens heading their colony. We then investigated whether queen acceptance was favored in specific conditions. We found that workers from polygynous colonies accepted alien queens when these queens were accompanied by workers. These results show that workers flexibly adjust their acceptance of alien queens according to the situation. We discuss how conditional acceptance of unrelated queens may be adaptive by providing benefits through increased colony size and/or genetic diversity, and by avoiding the rejection costs resulting from fighting.

## Background

Reproductive altruism has been a challenge for evolutionary theory for decades. How does a gene that makes an individual sacrifice or reduce its own reproduction spread in a population? This paradox was solved by W. D. Hamilton, who showed that alleles for altruism can increase in frequency when the recipient of altruism is genetically related to the actor, so that both have inherited identical alleles from recent common ancestors (Hamilton, 1963, 1964). An altruistic behavior will spread when the cost to the actor is smaller than the benefit to the recipient weighted by the level of genetic relatedness between them. This rule established relatedness as central for the evolution of reproductive altruism.

High relatedness pre-dates the evolution of cooperating cellular units (Fisher et al., 2013), of cooperative breeding in birds (Cornwallis et al., 2010), and of the sterile worker caste in eusocial species (Hughes et al., 2008). Yet despite relatedness being fundamental for the evolution of reproductive altruism, eusocial species have repeatedly evolved social structures that decrease relatedness between colony members. In about 44% of all ant species (Boomsma et al., 2014), colonies accept additional reproductive queens (a phenomenon called ‘secondary polygyny’, which can be facultative or obligate; hereafter referred to as ‘polygyny’). Although queens in polygynous nests are typically related to one another, colonies sometimes adopt unrelated queens (Stille and Stille, 1992; Seppä, 1996; Goodisman and Ross, 1999; Zinck et al., 2007). This phenomenon is puzzling as workers and resident queen(s) do not gain indirect fitness benefits through the reproduction of unrelated queens. Why then, would they accept these queens in their nest?

Cooperation among unrelated individuals should only evolve and be maintained when the fitness benefits exceed the costs for each partner. This can occur in ants if, for example, quickly increasing colony size provides a competitive advantage to all colony members. Accepting unrelated individuals may also be adaptive when, for example, a higher genetic diversity improves colonies’ resistance to parasites or increases their foraging efficiency (Schmid-Hempel and Crozier, 1999; Hughes and Boomsma, 2004; Mattila and Seeley, 2007; Seeley and Tarpy, 2007; Whitehorn et al., 2011). Direct benefits linked to increased group size and to higher genetic diversity are diverse and widespread (Avilés and Tufino, 1998; Bernasconi and Strassmann, 1999; Schmid-Hempel and Crozier, 1999; Mattila and Seeley, 2007; Donaldson-Matasci et al., 2013; Guindre-Parker and Rubenstein, 2020). Therefore, workers can gain indirect fitness benefits from accepting alien queens in their colonies if the presence of these unrelated queens increases the survival and reproductive success of queens to which they are related.

Alternatively, the acceptance of unrelated queens in eusocial colonies may result from failures in recognition systems of workers (Reeve, 1989; Vander Meer and Morel, 1998; Adams et al., 2007). Nestmate recognition in ants, as in other social insects, is based on the detection of non-volatile chemicals cues, mostly long chain hydrocarbons present on their cuticle (d’Ettorre and Lenoir, 2010; van Zweden and d’Ettorre, 2010). The exchange of cuticular compounds among colony members leads to the formation of a specific colony odor profile that allows individuals to discriminate nestmates from non-nestmates (Vander Meer and Morel, 1998; Hefetz, 2007; d’Ettorre and Lenoir, 2010; Sturgis and Gordon, 2012). Similarities in the odor profiles of colonies can lead to the erroneous acceptance of non-nestmate individuals (Reeve, 1989). Such errors might be more frequent in polygynous colonies, as the increased genetic diversity resulting from the presence of multiple matrilines in the colony may broaden the colony odor template (Reeve, 1989; Starks et al., 1998; Adams et al., 2007). In support of this view, workers from polygynous species are often less aggressive towards non-nestmate individuals than workers from monogynous species (i.e. species with a single reproductive queen) (Vander Meer and Morel, 1998). Moreover, aggressiveness towards non-nestmate individuals co-varies negatively with colony queen number in *Pseudomyrmex pullidus* (Starks et al., 1998), but see (Helanterä et al., 2011). Overall, workers from polygynous colonies might thus be more prone to accept non-nestmates queens, perhaps at the expense of their indirect fitness.

Here, we investigate whether the acceptance of unrelated queens by workers is incidental and results from a low ability of workers at recognizing alien queens, or whether it may be adaptive and favored in contexts where cooperation is preferable over rejection. Our model species is the socially polymorphic Alpine silver ant, *Formica selysi*. This species has two types of colonies co-existing within populations. Some colonies are headed by multiple queens, and others by a single queen. Social organization is determined by a supergene with two non-recombining haplotypes, Sm and Sp (Purcell et al., 2014; Avril et al., 2019). All single-queen colonies (hereafter denoted as monogynous colonies) are headed by a SmSm queen mated to one or two Sm males. Hence, all workers and gynes produced by monogynous colonies have the SmSm supergene genotype (hereafter monogynous workers and monogynous queens). Whereas all queens heading multiple-queen colonies (hereafter called polygynous colonies) have at least one copy of the Sp haplotype (Purcell et al., 2014; Avril et al., 2019), producing workers and gynes that have either the SmSp or SpSp supergene genotype (hereafter polygynous workers and polygynous queens). The relatedness between nestmate queens from polygynous colonies, although significantly higher than zero, is low and variable (r = 0.179 ± 0.018; mean ± SE; range = 0-0.9; (Avril et al., 2019)). Out of 179 pairs of nestmate queens, 67 (37%) were unrelated (r = 0), and 96 (53%) had relatedness estimates below 0.1 (pairwise relatedness estimates obtained from data in (Avril et al., 2019), using the method described in (Huang et al., 2015)). High relatedness between pairs of nestmate queens suggests that daughter-queens are re-accepted in their maternal nest, as occurs in many polygynous species (Bourke and Franks, 1995), whereas very low relatedness between pairs of nestmate queens suggests that adoption of non-nestmate, unrelated, queens occurs occasionally, as also happens in other polygynous species (Stille and Stille, 1992; Seppä, 1996; Goodisman and Ross, 1999; Zinck et al., 2007).

A previous study showed that *F. selysi* workers discriminate nestmate from non-nestmate workers, independently of whether they live in polygynous or monogynous colonies (Rosset et al., 2007). That study did not find any correlation between the level of aggression that workers displayed towards non-nestmate workers and the level of genetic diversity within their nest. Aggression levels were found to be higher between than within social forms. That study was carried out in the context of intraspecific colony competition and analyzed the behavior of workers towards alien workers. The behavior of workers towards alien queens is currently unknown.

In controlled behavioral experiments, we tested the mechanisms underlying the acceptance of non-nestmate queens by *F. selysi* workers. We focused on polygynous workers, as monogynous colonies typically do not host extra reproductive queens in nature (Purcell et al., 2014; Avril et al., 2019). We used relatively small colonies because, besides practical reasons, the potential benefits of accepting unrelated individuals may be higher in these colonies. In a first experiment we assessed whether the presence of unrelated queens in polygynous colonies could be incidental and result from lower discrimination abilities of polygynous workers. We considered two possibilities: 1) that polygynous workers (i.e. carrying the Sp haplotype) are inherently less capable at recognizing alien queens than monogynous workers (i.e. carrying the SmSm supergene genotype), and 2) that the presence of multiple queens in polygynous nests impairs the discrimination abilities of polygynous workers. Once we discovered that polygynous workers were as efficient as monogynous workers at recognizing and rejecting alien queens, independently of the number of reproductive queens heading their nest, we carried out a second experiment to investigate whether the acceptance of unrelated queens in polygynous colonies could be adaptive in some conditions. Specifically, we tested whether polygynous workers were more likely to accept unrelated queens that are accompanied by daughter workers than unrelated queens that are alone. Accepting queens with workers could be adaptive, particularly for small colonies, because it allows them to quickly increase colony size and colony genetic diversity (see above). Furthermore, rejecting alien individuals may be costly, especially when these individuals are numerous and defend themselves against their host. Therefore, we predicted that polygynous workers would be more likely to accept alien queens when these queens were accompanied by daughter-workers than when they were alone.

## General methods

**Experiment one: Are polygynous workers inherently less capable at recognizing alien queens? Is their ability to recognize alien queens reduced by having multiple queens in their nest?**

### Collection of host colonies

We collected fragments of mature, queenright colonies from a well-studied population in Finges, Valais, Switzerland (46.3138° N, 7.6012° E; 400m a.s.l.). *F. selysi* usually nests under stones. During spring, queens come to the top of their colonies to warm up and resume egg laying. We collected workers and queens from monogynous and polygynous colonies using tweezers, between March and May 2018. We used a SNPs genotyping qPCR assay to determine the supergene genotype of three workers per colony and infer the social form of their colonies (Foncuberta-De Gasperin et al., in prep). In each monogynous colony we collected the single queen, whereas in each polygynous colony we sampled one to eight queens. As some queens might have remained lower in the ground, the number of queens collected in polygynous colonies was probably below (but likely proportional to) the real number of queens heading these colonies. We placed each colony inside a plastic box (26 x 17 x 13.5cm), lined with fluon, and with a glass tube (length = 16cm; ø = 5mm) 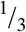 filled with water. We kept the colonies in the laboratory for 14 months before the start of the acceptance experiments, so that the sampled queens had time to produce daughter workers. Between May and October 2018, colonies were kept at 25° C, 70% humidity, in a light:dark 12:12 hours cycle, with food in the form of sugar and egg-jelly *ad libitum*. Between November 2018 and April 2019, the period of hibernation in nature, we kept colonies at 8° C, 70% humidity and in a light:dark 12:12 hours cycle, without food.

### Collection and experimental mating of alien queens

In June 2019, we collected sexual pupae and workers from another set of field colonies from the same population. We determined the social form of these colonies by using qPCR assay as described above. We kept the pupae and workers from each colony inside a plastic box (15.5 x 13.5 x 5.5cm), lined with fluon, and with a glass tube (length = 16cm; ø = 5mm) 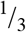 filled with water. We kept these colony samples in standard laboratory conditions, at 25° C, 70% humidity and in a light:dark 12:12 hours cycle, and fed them twice a week with food in the form of sugar and egg-jelly. Emerging queens and males were separated regularly to prevent them from mating. Young queens were placed to mate in artificial swarms alongside non-nestmate males, within plastic boxes (height = 20cm; length at the top = 42cm; width at the top = 26.5cm), covered with a mesh and placed under direct sunlight. We collected any mating pair and, four hours after mating, we introduced the queens into their host colony (see below).

### Introduction of alien queens into host colonies

At least two weeks before the experiment (in June 2019) we placed inside each colony a petri dish divided in two by a mesh and closed with a cover drilled with small holes. The set-up allowed workers to move freely in and out of the petri dish but prevented any queen from entering it, exiting it, or moving between the compartments inside the petri dish. At this point we counted all the workers and pupae of all colonies. Newly mated queens (see above) were introduced into three types of host colonies: i) monogynous colonies (n = 18); ii) single-queen polygynous colonies (n = 17); and iii) multiple-queen polygynous colonies (n = 18). Host colonies were relatively small (median size of 220, range = 57 – 884), and of similar sizes in the three conditions (Kruskal-Wallis χ^2^ = 3.10; d.f. = 2; *p =* 0.21). Polygynous colonies with multiple queens had a median number of 3 queens (range = 2 – 6).

We introduced one newly mated polygynous queen and one newly mated monogynous queen simultaneously into the petri-dish, randomly allocating each queen into one of the two compartments. Queens (alien and resident) were individualized via paint-marking before the trials. We monitored the survival of alien queens for 48hrs.

### Experiment two: Is queen acceptance by polygynous workers conditional and favored when alien queens are accompanied by workers?

We used 67 host colonies as described above. This time each replicate consisted of a pair of colonies (or a colony paired to a single queen), connected via a small foraging arena (plastic box of size 10.5 x 13.5 x 5.5cm) placed at equal distance of the two colonies. We randomly chose the side where each colony (or queen) was placed. Each colony was connected to the foraging arena through a rubber tube (ø = 5mm; length = 4cm). We had three treatments, with the focal colony being always a polygynous colony. The focal polygynous colony was connected to either: i) another polygynous colony (‘P-P’ treatment; n = 12), ii) a monogynous colony (‘P-M’ treatment; n = 15), or iii) a lone, newly mated, polygynous queen (‘P-SQ’ treatment; n = 12). Focal colonies headed by a single or by multiple queens were distributed evenly and randomly across all three treatments. To acquire newly mated, lone polygynous queens, we followed the same procedure as described for experiment one, and again introduced them into the experiment four hours after mating. Again, we individualized all queens by paint-marking them before the trials, at the same time for all queens. We predicted that polygynous colonies would kill lone polygynous queens, as in experiment one, but that they would accept polygynous queens with workers, leading to the merging of colonies, with non-nestmate queens sharing the same nest. We also predicted that workers would not accept monogynous queens, be it because of intolerance of monogynous and/or of polygynous workers (as monogynous colonies typically do not host extra reproductive queens and colonies hosting both queen types have not been found in nature, Purcell et al., 2014; Avril et al., 2019).

An observer naïve to the hypotheses and to the genotype of the queens and workers monitored the experiment twice a week, recording the status of each queen (dead or alive) and their spatial location (queens were allowed to move freely in this experiment). We replaced the food in the foraging arena twice a week. At the end of the experiment (after two months), we categorized the outcome of each replicate as acceptance if queens were observed sharing a nest the majority of times. In three replicates within the P-P treatment one of the queens died at the beginning of the experiment, but the remaining queens ended up living together with the queen(s) from the other colony. Whether we considered these three replicates as acceptance or not did not change the outcome of the statistical analyses. We consider these replicates as acceptance in the results presented below.

### Statistical analyses

All analyses were carried out in R (Team, 2014) v. 3.5.1. No statistical analysis was carried out for experiment one, as all alien queens were killed, in all treatments. For experiment two, we evaluated the prediction that polygynous colonies would be more likely to accept alien polygynous queens with workers than without workers. We coded ‘acceptance’ as one, and any other outcome (whether the queens died or remained alive but in separate nests) as 0 and ran a logistic regression (model 1). We included as explanatory variables the treatment (with three levels) and the number of queens in the focal colony. Because there was not a single replicate within the P-M treatment with an ‘acceptance’ outcome, we encountered complete separation (i.e. perfect correlation between the response and the treatment), detected by extremely large SE for each estimate. We thus ran a Firth’s bias-reduced logistic regression (package ‘logistf’ (Firth, 1993; Heinze et al., 2013)) and used the P-P treatment as a reference for comparisons with the P-SQ and P-M treatments.

We then used logistic regression to test whether queens had different probability to die during the experiment according to the treatment (model 2). If at least one of the queens died during the experiment (from the focal and/or non-focal colony), and the remaining queens did not end up sharing a nest, we coded the outcome as one, and the other outcomes as 0. Again, we included as explanatory variables the treatment and the number of queens of the focal colony. We obtained estimates with ANOVA type II SS (‘Anova’ function; (Fox et al., 2012)), and adjusted post-hoc *p*-values with FDR correction (‘lsmeans’ function; (Lenth, 2016)). As the number of queens heading the focal nests did not predict any response variable, we removed this from the minimal adequate models. Because the P-M and P-SQ treatments had the same statistical probability to have dead queens (from the focal and/or non-focal colony), we additionally tested whether the probability that the focal queen(s) would survive until the end of the experiment differed between treatments (model 3). Because no focal queen died within the P-SQ treatment (Figure 1), we again encountered complete separation and thus ran a Firth’s bias-reduced logistic regression, as above. We first used the P-M treatment as a reference and compared it with the P-P and P-SQ treatments, and then ran this model again excluding the ‘P-M’ treatment to evaluate differences between the P-P and P-SQ treatment.

**Figure 1.**
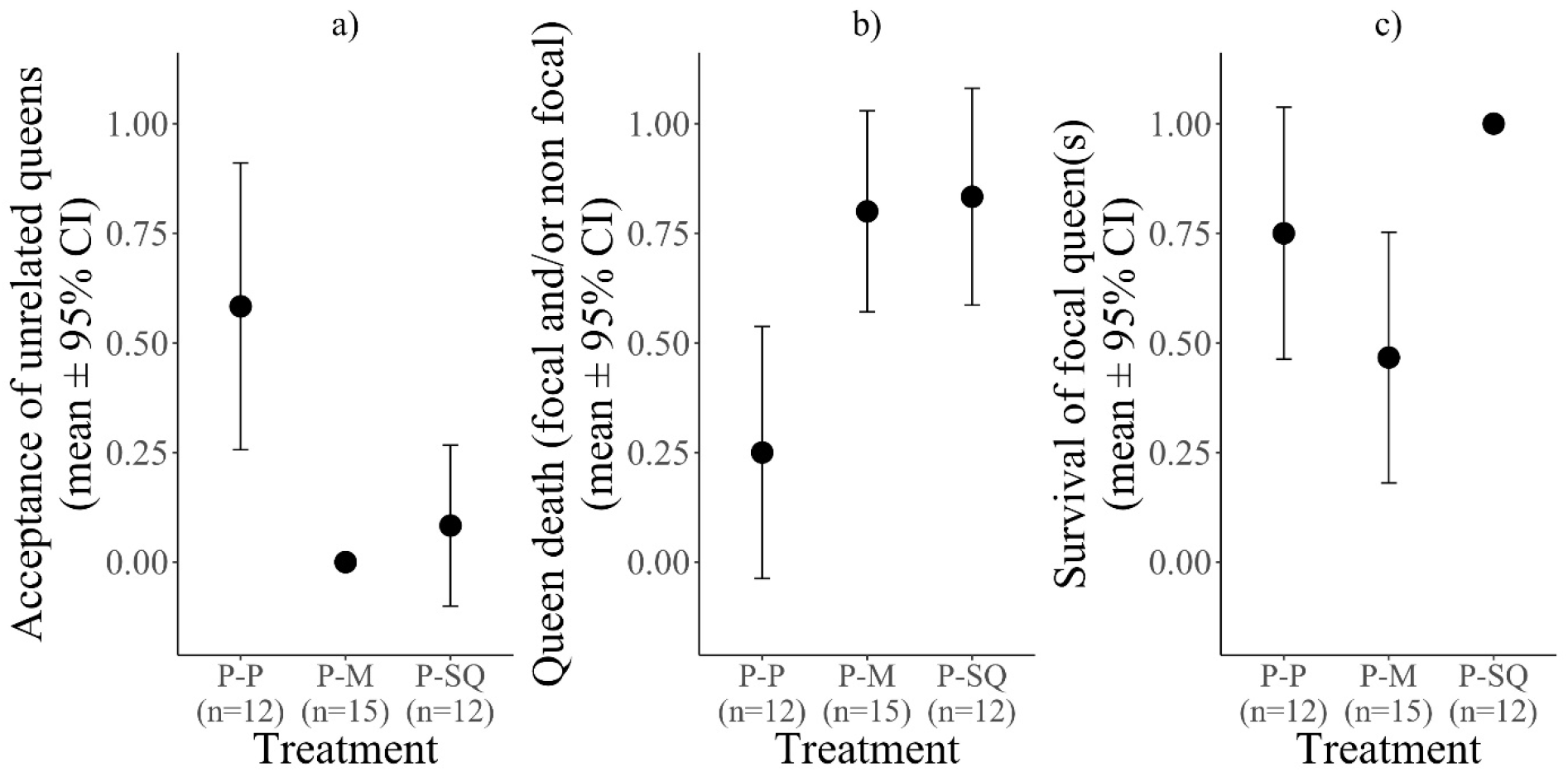
a) Probability of acceptance of alien queens by polygynous workers. Alien queens were considered to be accepted by the focal colony when focal and alien queens were seen a majority of time cohabiting in the same nest; b) proportion of replicates with queen death. The outcome of the replicate was considered ‘queen death’ when at least one queen from the focal or non-focal colony died, and when the focal and non-focal queens left alive did not share a nest the majority of times; and c) proportion of replicates where the focal queen(s) survived until the end of the experiment. All graphs show the mean ± 95% CI, per treatment. All results are from experiment

## Results

### Experiment one: Are polygynous workers inherently less capable at recognizing alien queens? Is their ability to recognize alien queens reduced by having multiple queens in their nest?

Polygynous workers were as efficient as monogynous workers at recognizing and eliminating alien queens. All alien queens were killed within 48hr. Hence, rejection occurred irrespective of the social origin of the introduced alien queens, the genotype of the workers, or the number of queens in the host colony (number of host colonies: polygynous with multiple queens n = 17; polygynous with a single queen n = 18; monogynous = 18).

### Experiment two: Is queen acceptance by polygynous workers conditional and favored when alien queens are accompanied by workers?

As predicted, polygynous workers were more likely to accept alien polygynous queens when these queens were accompanied by workers than when they were alone (model 1; acceptance in the P-P vs. P-SQ treatments: Coef = -2.35 with 95% CI [-4.78, -0.51]; χ^2^ = 6.50; *p* = 0.01; Figure 1a). Acceptance rate of alien queens accompanied by workers was higher when these queens and workers belonged to the polygynous form than when they belonged to the monogynous form, also as predicted (model 1; acceptance in the P-P vs. the P-M treatment: Coef = -3.74 with 95% CI [-8.67, -1.39]; χ^2^ = 11.99; *p* < 0.001; Figure 1a).

Queen death (in the focal and/or non-focal colony) occurred proportionally less often when both the focal and non-focal colonies had polygynous queens with workers (model 2; overall χ^2^ = 11.59: *p* = 0.003; Figure 1b). The focal queen(s) had similar probabilities of surviving until the end of the experiment in the P-P and in the P-M treatment (model 3; P-P and P-M comparison: coef = 1.12 with 95% CI [-0.39, 2.80]; χ^2^ = 2.07; *p* = 0.15), but, at the replicate level, death of queen (either focal or non-focal) was more probable in the P-M than in the P-P treatment (model 2; *post-hoc* comparison: Estimate = 2.48; SE = 0.92; *z* = 2.67; *p* = 0.01; Figure 1b). Queens were thus more likely to die when monogynous colonies were involved, than when only polygynous colonies were involved. Queen death was also more likely in the P-SQ than in the P-P treatment (model 2; *post-hoc* comparison: Estimate = -2.70; SE = 1.02; *z* = -2.65; *p* = 0.01; Figure 1b) and focal queens had marginally larger survival probabilities in the former than in the latter treatment (model 3; P-P and P-SQ comparison: coef = 2.22 with 95% CI [-0.27, 7.16]; χ^2^ = 2.94; *p* = 0.08). The probabilities that some queens died were similar in the P-M and P-SQ treatments (model 2; *post-hoc* comparison: Estimate = -0.22; SE = 1.00; *z* = -0.22; *p* = 0.82). However, only the newly-mated polygynous queen died in the P-SQ treatment (Figure 1c), whereas it was often the focal polygynous queen(s) who died in the P-M treatment (model 3; P-SQ and P-M comparison: coef = 3.34 with 95% CI [1.03, 8.25]; χ^2^ = 9.56; *p* = 0.001; Figure 1c). The number of queens within the focal colony did not correlate with probabilities of queen acceptance, queen death, or survival of focal queens (all *p* > 0.50).

## Discussion

Kin selection solved one of the greatest mysteries in evolutionary biology: the evolution of reproductive altruism. When a gene makes an organism help close kin reproduce, it is passing on copies of itself to future generations, not through direct descent, but through copies of itself present in others. High relatedness predates the evolution of multicellularity (Fisher et al., 2013), of cooperative breeding (Cornwallis et al., 2010), and of eusociality (Hughes et al., 2008). Yet almost half of all known ant species have colonies with multiple reproductive queens (Boomsma et al., 2014), which can sometimes be unrelated (Stille and Stille, 1992; Seppä, 1996; Goodisman and Ross, 1999; Zinck et al., 2007)). Why do workers accept unrelated queens in their nest, when they do not gain indirect fitness through their reproduction? Our results help answer this question by showing that the acceptance of unrelated queens can be context-dependent and likely happens when the benefits of accepting them outweigh the costs (see (Sturgis and Gordon, 2012)). Workers from polygynous colonies accepted alien queens, but only when these queens were accompanied by daughter-workers. Lone queens were killed within hours.

We suggest that this constitutes a form of mutualism, where individuals from both colonies benefit from the association. Increasing colony size may increase colony survival, because larger colonies have improved nest thermoregulation (Korb, 2003; Jones and Oldroyd, 2006; Kadochová and Frouz, 2013), foraging efficiency (Donaldson-Matasci et al., 2013), division of labor (Holbrook et al., 2011; Ferguson-Gow et al., 2014), and nest defense abilities against competitors and predators (Guindre-Parker and Rubenstein, 2020). Moreover, having genetically diverse, albeit unrelated, individuals within the nest may also enhance inclusive fitness of all colony members, as increased genetic diversity may lead to better colony immunity (Schmid-Hempel and Crozier, 1999; Hughes and Boomsma, 2004; Seeley and Tarpy, 2007), colony homeostasis (Oldroyd and Fewell, 2007) and foraging efficiency (Mattila and Seeley, 2007). Therefore, the integration of numerous unrelated individuals into a colony might bring ecological benefits to members of both colonies. Additionally, or alternatively, tolerating alien queens when accompanied by workers may be the ‘best of a bad job’, where peaceful cooperation is less costly than losing workforce and queens during fights with other colonies (as happened when encountering monogynous colonies). Whatever the relative importance of these two mechanisms, we suggest that individuals gained stronger net benefits when cooperating with alien queens and workers than when rejecting them, hence making acceptation of unrelated queens probably adaptive in this context.

The potential benefits of increasing colony size may have been particularly large for our experimental colony fragments, due to their relatively small size compared to field colonies (mature polygynous colonies have on average 30,000 workers (Rosset and Chapuisat, 2007)). Indeed, the benefits of increased colony size may not be linear and rather follow an inverted U shape, reaching an upper-limit and then decreasing beyond a certain colony size, as large colony size may lead to food depletion (Bonal and M. Aparicio, 2008), inefficiencies in task performance (Michener, 1964) and/or reduced worker longevity (Rueppell et al., 2009; Blacher et al., 2017). Investigating the importance of colony size (both in absolute terms and relative to the size of neighboring colonies) in the probability of accepting alien queens and their workers are promising lines of research for future work.

Polygynous workers were as efficient as monogynous workers at detecting and killing alien queens, independently of the number of queens heading their colony. Because recognition cues have a heritable component (Gamboa et al., 1986; Adams, 1991; van Zweden et al., 2010), having many matrilines in a colony increases the diversity of recognition cues within the nest. This could in turn affect nestmate recognition mechanisms, generating a positive feedback between the number of reproductive queens in the colony and the acceptance of non-nestmates. Yet we did not find evidence of such a feedback in our experiments, as the number of queens heading the nests did not correlate with the acceptance rate of alien queens, in any of the two experiments. This is also supported by a previous experiment in the field, which found that workers living in polygynous colonies recognize nest-mate workers as effectively as those living in monogynous colonies (Rosset et al., 2007). The presence of unrelated queens in natural polygynous colonies of *F. selysi* (Avril et al., 2019) is hence unlikely to be generally explained by an inherently lower ability of polygynous workers at distinguishing non-nestmate queens, or by a more permissive acceptance threshold in polygynous workers caused by a wider diversity of odor cues in their nest.

To conclude, our experiment helps explain why unrelated reproductive queens are occasionally present in eusocial insect colonies (Stille and Stille, 1992; Bourke and Franks, 1995; Seppä, 1996; Goodisman and Ross, 1999; Zinck et al., 2007; Boomsma et al., 2014), despite the fact that colony members do not gain indirect fitness from their reproduction. Polygynous workers rejected lone alien queens, but frequently accepted them when these queens were accompanied by workers. The acceptance of unrelated queens was context-dependent (Sturgis and Gordon, 2012) and probably occurred when the direct benefits of acceptance, in terms of colony growth, survival and productivity, outweighed the costs of rejection. Our study hence conflicts the common view that the presence of alien, unrelated, individuals in colonies of social insects is solely due to failures in their recognition system (see Reeve, 1989; Vander Meer and Morel, 1998; Adams et al., 2007; Fouks et al., 2011). We propose that unrelated queen acceptance can be adaptive in specific contexts where peaceful cooperation is preferable to risky rejection.

## Acknowledgments

We thank Sagane Dind, Solenn Sarton-Lohéac, Marjorie Labédan, Jason Buser and Christophe Lakatos for their help with laboratory work.

